# The rt-TEP tool: real-time visualization of TMS-Evoked Potential to maximize cortical activation and minimize artifacts

**DOI:** 10.1101/2021.09.15.460488

**Authors:** S Casarotto, M Fecchio, M Rosanova, G Varone, S D’Ambrosio, S Sarasso, A Pigorini, S Russo, A Comanducci, RJ Ilmoniemi, M Massimini

## Abstract

**Background:** The impact of transcranial magnetic stimulation (TMS) on cortical neurons is currently hard to predict based on a priori biophysical and anatomical knowledge alone. This problem can hamper the reliability and reproducibility of protocols aimed at measuring electroencephalographic (EEG) responses to TMS.

**New Method:** We introduce and release a novel software tool to facilitate and standardize the acquisition of TMS-evoked potentials (TEPs). The tool, `rt-TEP` (real-time TEP), interfaces with different EEG amplifiers and offers a series of informative visualization modes to assess in real time the immediate impact of TMS on the underlying neuronal circuits.

**Results:** We show that `rt-TEP` can be used to abolish or minimize magnetic and muscle artifacts contaminating the post-stimulus period thus affording a clear visualization and quantification of the amplitude of the early (<50 ms) EEG response after averaging a limited number of trials. This real-time readout can then be used to adjust TMS parameters (e.g. site, orientation, intensity) and experimental settings (e.g. loudness and/or spectral features of the noise masking) to ultimately maximize direct cortical effects over the undesired sensory effects of the coil’s discharge.

**Comparison with Existing Methods:** The ensemble of real-time visualization modes of `rt-TEP` are not implemented in any current commercial software and provide a key readout to titrate TMS parameters beyond the a priori information provided by anatomical models.

**Conclusions:** Real-time optimization of stimulation parameters with `rt-TEP` can facilitate the acquisition of reliable TEPs with a high signal-to-noise ratio and improve the standardization and reproducibility of data collection across laboratories.

**Highlights:**

- `rt-TEP` provides an immediate EEG readout to assess the quality of TEPs in real time
- `rt-TEP` interfaces with most commercial EEG systems
- Informative, real time visualization allows maximising the cortical impact of TMS while minimizing artifacts
- `rt-TEP` facilitates the acquisition of reliable TEPs with a high signal-to-noise ratio

## 1. Introduction

Transcranial magnetic stimulation (TMS) can directly activate the cerebral cortex with a huge number of parameter combinations such as position, orientation, and intensity of the induced electric field (E-field). This flexibility offers unprecedented opportunities for exploring and modulating cortical excitability but also represents a challenge; when a TMS probe is positioned on the scalp region overlying a cortical area of interest, the actual impact of the E-field on cortical neurons is very hard to predict. Indeed, even when individual head models provided by state-of-the-art TMS navigation systems are available as a priori information, key factors such as microscale axon orientation, cytoarchitectonics and local neuronal excitability remain unaccounted for and may dramatically affect the interaction between the induced E-field and brain activity.

For this reason, when targeting the primary motor cortex (M1), TMS stimulation parameters (position, angle, intensity) are initially set based on coarse a priori anatomical information (Silva et al., 2020) and then adjusted post-hoc, until the electromyographic (EMG) activity recorded from a selected target muscle satisfies standard latency and amplitude requirements (Rossini et al., 2015). Hence, despite the potential confound represented by changes in the excitability of spinal motor neurons, EMG waves as a real-time feedback have been the fundamental guide for titrating TMS in research, diagnostic and treatment protocols.

Unfortunately, when stimulating outside the primary motor cortex such immediate readout is not available, thus preventing a reliable control over whether, or how effectively, TMS is impacting cortical neurons. This lack of control represents a fundamental limitation not only for TMS protocols aimed at inducing plasticity but also for studies in which TMS is employed in combination with other techniques to assess the state of cortical circuits. For example, differences in the effectiveness of direct cortical activation have been highlighted as a major problem affecting the reproducibility of studies employing TMS in combination with electroencephalography (TMS–EEG) to probe cortical excitability and connectivity (Belardinelli et al., 2019). Clearly, maximizing the direct impact of stimulation on cortical neurons while minimizing collateral effects such as scalp muscle, magnetic or sensory activations is a key prerequisite for improving the reproducibility across laboratories, the signal-to-noise ratio (SNR) and the informativeness of TMS–EEG studies. Towards this aim, in our previous TMS–EEG works, we have used visualization software to set stimulation parameters in real time based on the quality and amplitude of the EEG response to TMS (Casali et al., 2010; Casarotto et al., 2016; Rosanova, Fecchio et al., 2018). Such software tools were either implemented on amplifiers that are now out of production (eXimia EEG, Nexstim Plc, Finland) or based on hardware/software solutions that were customized to our specific set-up and are thus not available to the larger community.

Here, we present `rt-TEP` (real-time TMS-Evoked Potential), an open-source, Matlab®-based software tool that allows quantifying in real time the direct impact of TMS on any cortical area while minimizing common confounds which include magnetic artifacts as well as muscle and sensory-related activations. By representing the early (0–50 ms) EEG response to TMS in informative, interactive displays, `rt-TEP` guides the operator in setting optimal stimulation parameters during the experiment while continuous raw data are stored on disk for subsequent data pre-processing and analysis. Using `rt-TEP`, the operator can (*i*) effectively inspect single-trial data to detect artifacts and minimize them by small adjustments of coil position and rotation and (*ii*) visualize in real time the amplitude of the early TEP average (e.g., 10-30 trials) and (*iii*) estimate its SNR and overall quality. We show how this real-time readout about the immediate EEG effects of TMS enables the selection of optimal stimulation parameters (e.g., site, orientation, intensity) and of other relevant experimental settings (e.g., loudness and/or spectral features of the noise masking; see Russo et al., 2021) in order to maximize the strength of direct cortical activation over noise, artifacts and other undesired effects of the coil’s discharge. `rt-TEP` interfaces with the most widely used TMS-compatible EEG amplifiers. Source code is released (https://github.com/iTCf/rt-TEP.git) under GNU General Public License v3.0, which allows the user to extend the compatibility of this software to other EEG amplifiers.

## 2. Real-time control of TEPs with rt-TEP: rationale and procedures

`rt-TEP` software must be installed on a client computer (Windows, Mac or Linux OS) that receives real-time data streaming via ethernet from a server computer, which is in turn directly connected to the EEG amplifier and runs a proprietary software responsible for data collection and storage. As a first step, `rt-TEP` guides the operator through a series of interactive windows to select the EEG amplifier in use (**Supplementary Figure 1A**). Currently, four different amplifiers can be selected: BrainAmp (Brain Products GmbH, Germany), g.HIamp (G.TEC Medical Engineering GmbH, Austria), eego™ mylab (ANT Neuro, Netherlands) and Bittium NeurOne™ Tesla (Bittium Corporation, Finland).

Then, `rt-TEP` requires one to specify the settings for connecting with the server computer (e.g., IP address) and for desired data display (e.g., layout of the recording channels, temporal windows of interest, sampling rate) (**Supplementary Figure 1B**). In principle, any arbitrary set of EEG and EOG (electrooculographic) recording channels can be specified and visualized (**Supplementary Figure 1C**); additional channels connected to the same amplifier, such as electrocardiographic channels, must be included in the recording channels layout although are not displayed by `rt-TEP`.

Once connected to the amplifier, `rt-TEP` automatically displays continuous raw EEG data (see arrow 1 in **Figure 1**) to check for successful real-time data streaming. At this point, `rt-TEP` is ready to check TEP features. This can be done by activating two different interactive visualization modes (see arrow 2 and arrow 3 in **Figure 1**), updated in real time after each TMS pulse. Used sequentially, these modes guide the operator through a series of steps to optimize stimulation parameters before starting the actual measurement. The first mode allows an informative inspection of single-trial responses that is instrumental to (*i*) mask the pulse artifact, (*ii*) compute the average reference after the rejection of bad channels, (*iii*) assess and remove the recharge artifact, (*iv*) detect, locate and correct the discharge artifact, and (*v*) detect, localize and avoid/minimize the muscle artifact. Once all the artifacts potentially affecting the early post-stimulus time interval have been identified and minimized by the user, the second mode, which displays average data, offers a quantitative evaluation of the impact of TMS on cortical neurons. This is the critical step in which the operator can set the final stimulation parameters to ensure that the early EEG response to TMS is present and falls within the desired amplitude range, and that no obvious sensory-related artifacts are present. In typical conditions, this EEG-guided parameter search lasts for about 10 min from the initial coil positioning on the area of interest. In what follows, we illustrate this process and explain its rationale by using practical examples and real data.

**Figure 1.**
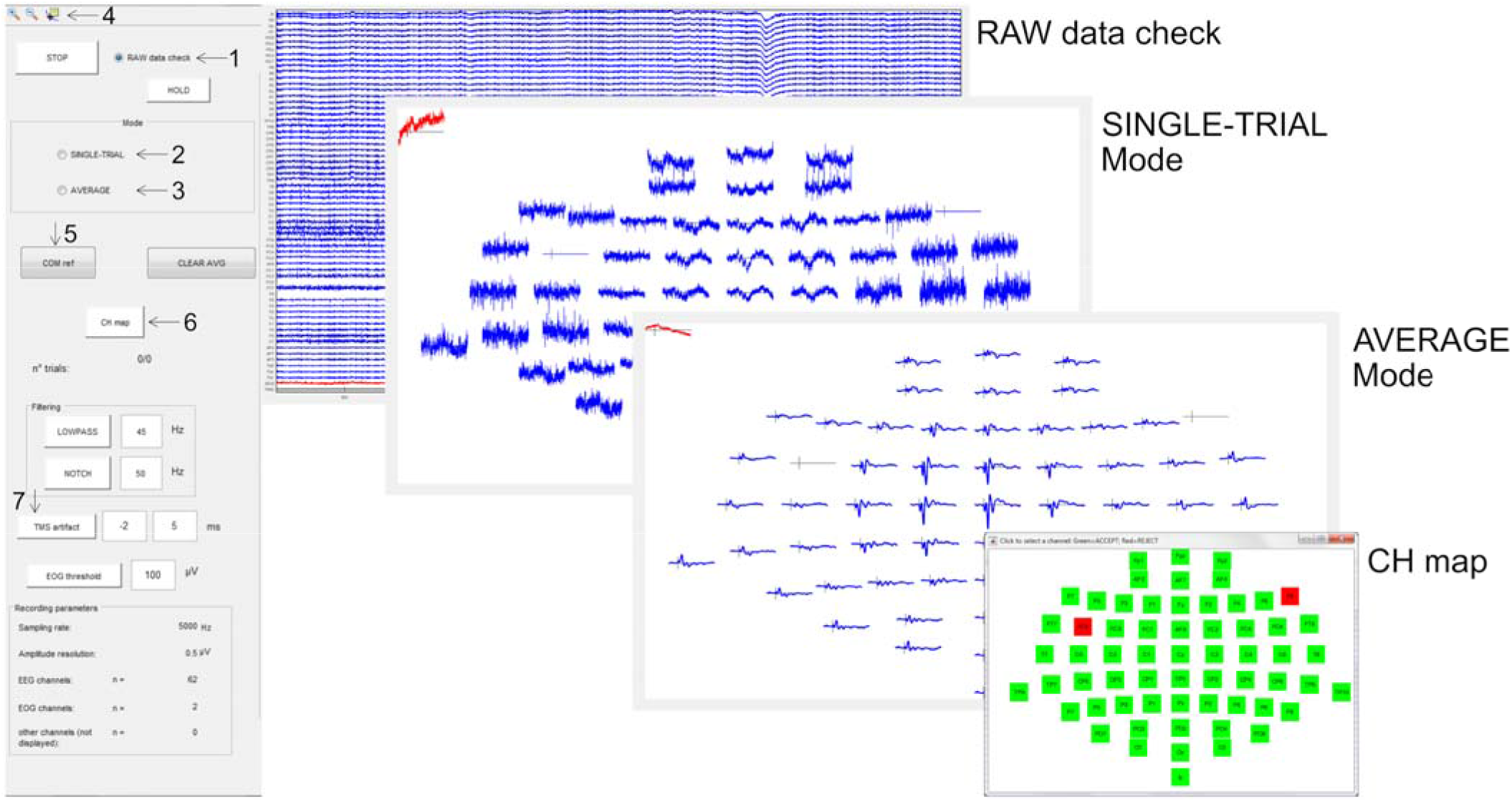
The main control panel of `rt-TEP` (left) and the three different displays that can be alternatively activated, including RAW data check, SINGLE-TRIAL Mode and AVERAGE Mode. In the topographic arrangement view, data are displayed in average reference by default; however, common physical reference can be retrieved by pressing the button COM ref. Zooming and measuring tools can be activated through the buttons on the top left menu bar (arrow 4). Button CH map opens an interface (shown in the bottom right corner) that allows to individually reject single-channel data from the display as well as from the computation of the average reference. When the TMS artifact button is activated, a selectable time window around TMS pulses is flattened to mask the pulse artifact. In addition, `rt-TEP` can apply LOWPASS and NOTCH filtering in real time to both single-trial and average data; when the EOG threshold button is activated, single trials with an EOG amplitude exceeding a selectable value are automatically excluded from the averaging process.

### 2.1 SINGLE-TRIAL mode: minimization of short-latency artifacts

In single-trial mode, `rt-TEP` shows EEG epochs of specified duration around the TMS pulse, refreshing at every stimulus: EEG channels are displayed in a topographic arrangement and signals can be easily explored through simple zooming and measuring tools (see arrow 4 in **Figure 1**). In this mode, the user is able to quickly detect and avoid artifacts that contaminate the initial portion of the post-stimulus EEG, as illustrated in detail below.

#### 2.1.1 Pulse artifact

During TMS pulse delivery, the transient (a few hundreds of μs) flow of current in the coil, peaking at several kA (Koponen et al., 2015), induces a large voltage in the EEG leads, which is proportional to the time rate of the magnetic flux threading the loop formed by the positive and negative input of the EEG amplifier, along the wires to the electrodes and via the head. Typically, a customized electrode shape is employed to prevent the induction of eddy currents in the electrodes themselves (e.g., pellet-wise, wire-wise, ring-wise with a slit) (Virtanen et al., Med Biol Eng Comput 1999). The pulse artifact’s duration can be further reduced (down to a few milliseconds) by setting optimal acquisition parameters of the EEG amplifier, such as wide measuring range (tens of mV), a large hardware filtering bandwidth (up to several kHz) and a high sampling rate (several kHz). Even when these procedures are in place, the electromagnetic artifact is still several orders of magnitude larger than brain activity and hampers the real-time assessment of the early EEG response to TMS (**Supplementary Figure 2**).

As a first step, `rt-TEP` allows one to remove the large pulse artifact from the online visualization by replacing the real signal with a constant value within a selectable time window around pulse delivery (see arrow 7 in **Figure 1**). As an example, using the BrainAmp DC amplifier (Brain Products GmbH, Germany) with 16.384 mV measuring range (16 bit A/D converter, 0.5 μV resolution per bit), DC-to-1-kHz hardware filtering bandwidth and 5-kHz sampling rate, the pulse artifact can be effectively removed by setting the time window to be flattened from −2 to +5 ms with respect to the pulse (**Figure 2A,B**). Once the large pulse artifact is eliminated, the *y*-axis range can be uniformly re-scaled to proceed with a more fine-scaled inspection of the EEG signal.

**Figure 2.**
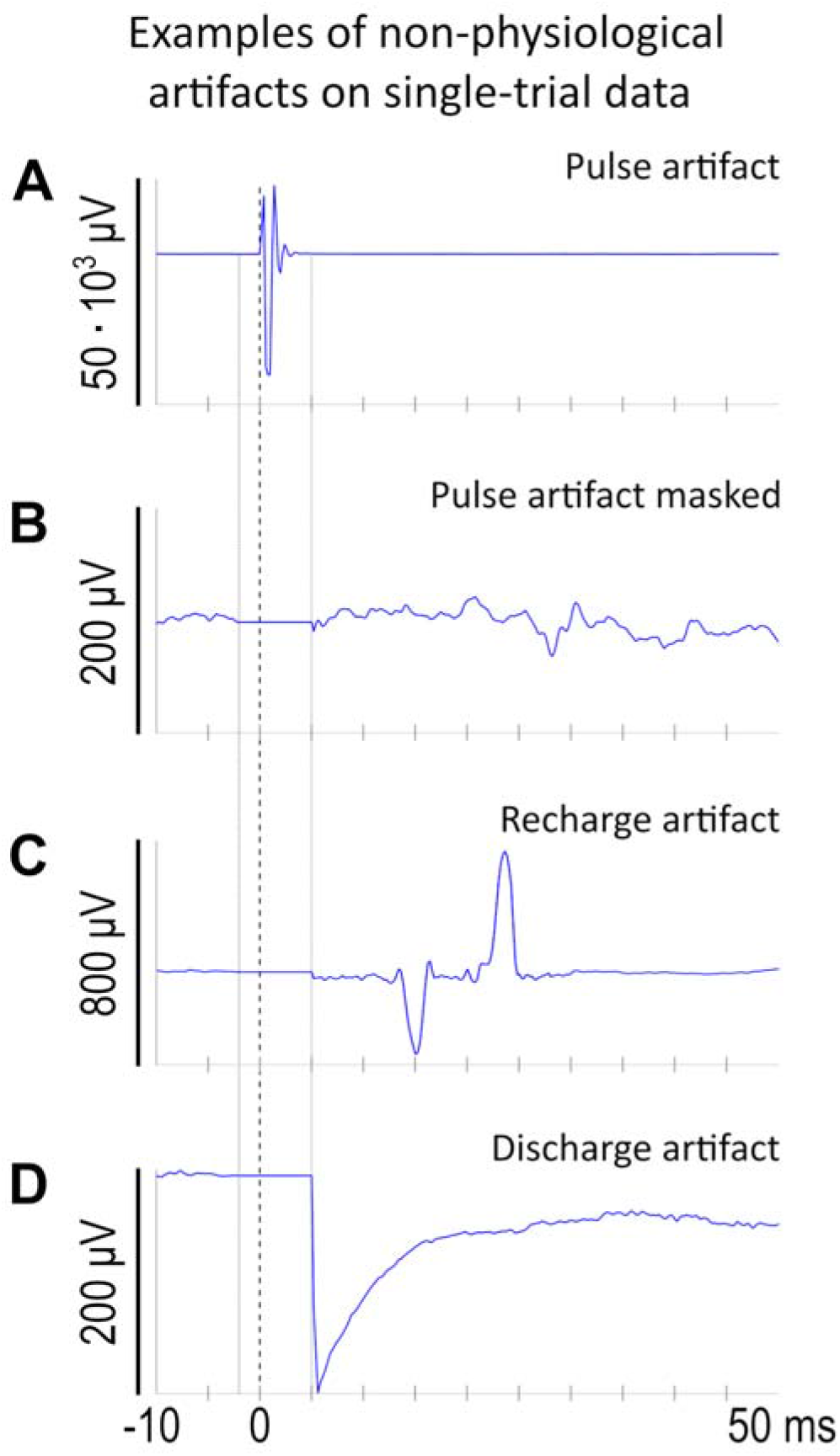
Examples of non physiological artifacts visible on single-channel, single-trial data: pulse artifact in common physical reference before (A) and after (B) masking, recharge artifact in average reference (C) and discharge artifact in average reference (D). For the all-channels display, see **Supplementary Figure 2**, **Supplementary Figure 3** and **Supplementary Figure 4**.

#### 2.1.2 From common physical reference to average reference: rejecting bad channels

The data streamed out by the amplifiers are obtained as potential differences between each measuring electrode and a unique “reference” electrode. These unipolar EEG recordings are inevitably affected by the location of the physical reference due to the lack of a “neutral” reference site anywhere in the body (Lei and Liao 2017; Nunez and Srinivasan 2006). Hence, the time course of either single-trial or average data in common reference results in an artificially high degree of correlation across channels. This prevents the detection of artifacts with a characteristic spatial distribution, the discrimination between widespread artifacts and common mode signals, and ultimately the evaluation of the topography of genuine EEG responses to TMS. In principle, the average reference, which consists in subtracting from each EEG recording a linear combination of the recordings from all electrodes, successfully mitigates the bias of unipolar reference, provides better spatial information and a more reliable estimation of the signal amplitude (Nunez, 2010; Yao et al., 2019). However, a reliable average reference montage requires that bad channels that are either saturated, disconnected, or contaminated by irreducible artifacts, are not considered; otherwise, artifacts would naturally spread across all the recording electrodes and/or the signal would be artificially injected into channels that are not picking up any electrical activity either because they are disconnected or saturated. In order to compute a reliable average reference, `rt-TEP` implements a simple interface to reject bad channels in real time (see arrow 6 in **Figure 1**): rejected channels are excluded from the computation of the average reference and are not displayed. The detection of bad channels can be performed both in average reference and in common reference, since `rt-TEP` allows to switch between these two montages with just a button press (see arrow 5 in **Figure 1**). This is important because if abnormal activity in some channels has such a large amplitude to contaminate all the other channels when computing the average reference, the detection of bad channels can be better performed by switching to common reference. For example, this kind of montage allows detecting outlier channels that are either heavily contaminated by artifacts (e.g., because of a loose contact) or flat (e.g., due to amplifier saturation). Such flexibility in the exploration and rejection of bad channels is not available in typical commercial acquisition software and is a prerequisite for the following steps as it allows visualizing average-reference EEG potentials on-line with unprecedented clarity.

#### 2.1.3 Recharge artifact

After zero-padding the pulse artifact and re-referencing, the third step involves inspecting the signal for other sources of artifact such as the recharge artifact. Indeed, EEG amplifiers may be also disturbed by the recharging of the stimulator’s capacitors between subsequent pulses; this occurs when there is a transient current flow or a change in the potential of the coil during the recharging. The occurrence of this artifact can be readily identified on the single-trial display provided by `rt-TEP`, once the pulse artifact is eliminated (**Supplementary Figure 3**). The waveform of this artifact may change with the characteristics of the TMS unit, the coil type, and the stimulation intensity. **Figure 2C** shows an example of the recharge artifact produced by the Rapid^2^ stimulator (Magstim Ltd, UK) equipped with a D70 Remote Coil. By default, the stimulator’s capacitors are usually recharged immediately after pulse delivery, thus possibly contaminating early EEG responses to TMS. If not detected before acquisition, the presence of a recharge artifact within a time-window of interest can irreversibly compromise the measurement.

If present, the recharge artifact can be avoided by delaying via software on the TMS unit the time of recharging beyond the temporal window of interest for TMS-evoked potentials. As an example, when stimulating at an inter-pulse interval randomly jittered between 2 and 2.3 s, the operator could set the time of recharging between 900 and 1000 ms after the pulse in order to get the largest symmetric artifact-free temporal window around TMS.

#### 2.1.4 Discharge artifact

Electromagnetic pulses may also charge stray or material-boundary capacitances located along any possible induction current path, e.g., at the electrode–gel and gel–skin interfaces. In particular, the area covered by the conductive gel and the aqueous ionic extracellular space of deep skin layers, separated by the stratum corneum of the epidermis acting as a hydrophobic dielectric, forms a capacitor (Freche et al., 2018). Immediately after being charged by the pulse, these capacitances discharge and produce a non-exponentially decaying artifact on the recorded signal, which can last several tens of ms. **Supplementary Figure 4** shows how this artifact appears on the single-trial display provided by `rt-TEP` after masking the pulse artifact (**Figure 2D** displays a representative channel).

The discharge artifact can be prevented by carefully lowering the resistance of the outermost layer of the epidermis and by using EEG electrodes with a shape that has been specifically designed to facilitate skin preparation. In particular, it is important (*i*) to move the hair from the scalp surface in direct contact with the electrode, (*ii*) to scrub the skin with an abrasive paste and (*iii*) to finally inject as much gel as necessary between the skin and the electrode. In case a decay artifact is still present, a further refinement of electrode impedance may be helpful in reducing its amplitude.

#### 2.1.5 Muscle artifact

The interaction between TMS pulses and excitable scalp tissues, such as muscular fibers and peripheral nerves, may produce additional artifacts that are superimposed on the brain responses due to direct perturbation of cortical neurons. Indeed, TMS pulses may induce scalp muscles to twitch by either the direct stimulation of their fibers or by the activation of their nerves. The resulting electromyographic signals, recorded by the EEG electrodes, can be several orders of magnitude larger than neuronal signals and can last tens of ms. Once the pulse, recharge and discharge artifacts are eliminated, `rt-TEP` enables a straightforward detection of muscle artifacts. Their waveform is usually characterized by a biphasic deflection, peaking at 5–10 ms and at 8–20 ms poststimulus, followed by a slow return to the baseline level at around 40–50 ms (Mutanen et al., 2013; Mäki and Ilmoniemi, 2011). Since scalp muscles are mostly located over the frontolateral and occipital aspects of the head, EEG responses to TMS are less susceptible to muscle artifacts when pulses are delivered over dorsal regions close to the midline (Mutanen et al., 2013).

When targeting sites located away from the medial aspect of the scalp, the likelihood of supra-threshold activation of scalp muscles by TMS depends on the stimulation intensity and on the angle between the main direction of muscle fibers and of the induced E-field. **Figure 3** shows two representative single-trial EEG responses to TMS (zoom on frontocentral channels; for the all-channels display see **Supplementary Figure 5**) obtained by stimulating the same cortical site, at the same intensity, but with a different orientation of the coil: the muscle artifact clearly visible in **panel A** is effectively abolished by simply rotating the coil 30° clockwise (**panel B**). This example shows that muscular artifacts can be successfully reduced by applying an EEG-informed fine-tuning of TMS parameters, which does not necessarily imply a major change of the target site or stimulation intensity. Minimizing muscle twitches before starting the measurements is very important for at least two reasons. First, it allows visualizing the early components (within the first 50 ms) of the EEG response to TMS, which are key for titrating the final stimulation parameters (see paragraph 2.2). Second, because they are clearly perceived by the examined subject, thus potentially resulting in unspecific brain responses to sensory stimulation (Conde et al., 2019).

**Figure 3.**
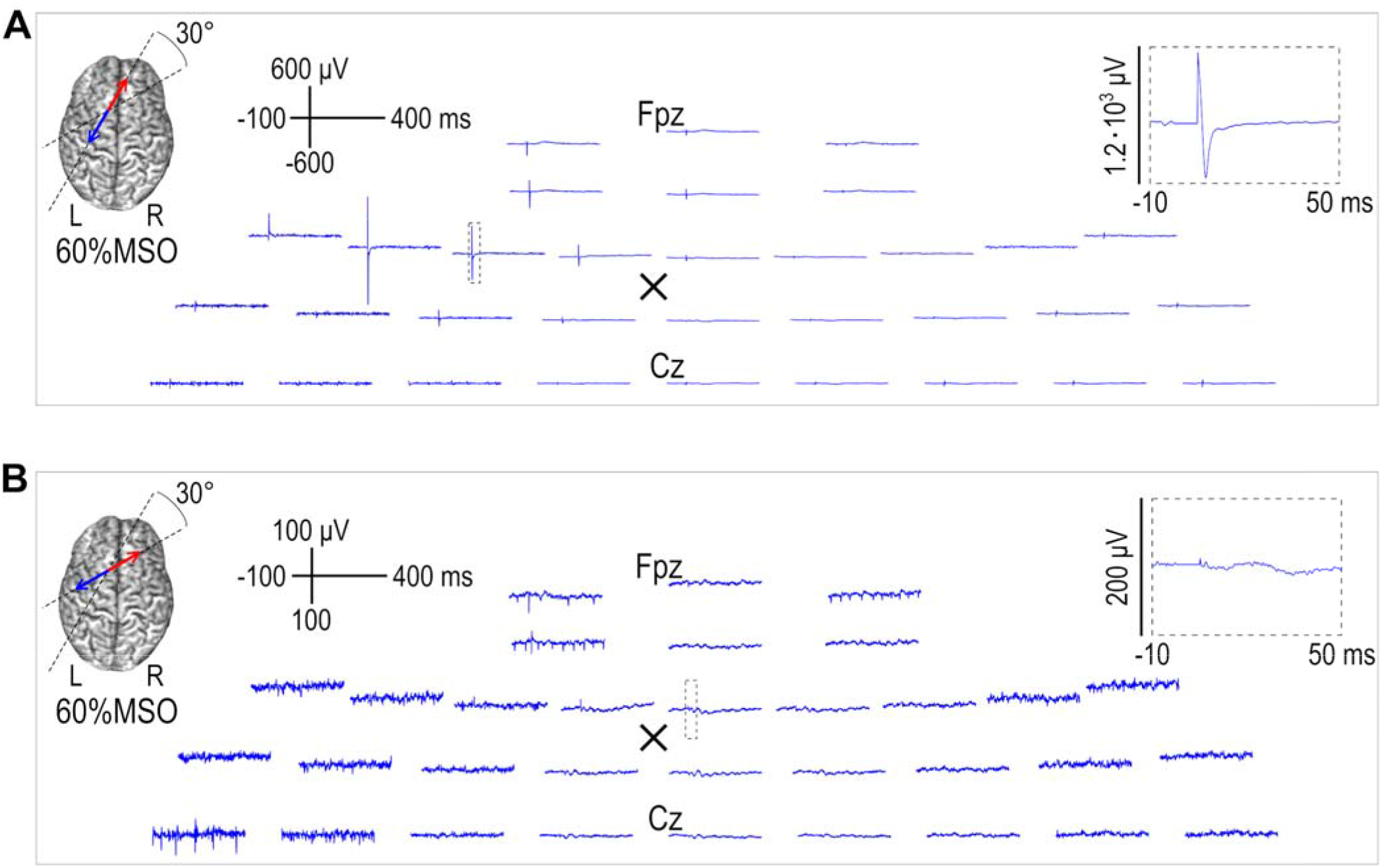
Single-trial EEG responses to TMS applied on the left frontal cortex (black cross) at 60%MSO (zoom on frontocentral channels): the orientation of the induced E-field differs by 30° between panels A and B, as depicted on a rendered brain. The pulse artifact has been visually masked by flattening the signal between −2 and +5 ms. EEG data were collected with a BrainAmp DC amplifier and TMS was delivered with an NBS4 Nexstim system. MSO = maximum stimulator output; L = left; R = right.

In conclusion, different artifacts can mask the initial brain responses to direct cortical stimulation with TMS. The single-trial display mode provided by `rt-TEP` guides the operator through a series of steps in which these artifacts can be recognized and eliminated. Once this is done, the TMS operator can move to the average display mode to visualize, quantify, and optimize the effects of TMS on the underlying circuits, before performing the actual measurement.

### 2.2 AVERAGE MODE: maximizing the signal-to-noise ratio of TEPs

In the average mode, `rt-TEP` provides real-time feedback about the amplitude, morphology and topography of the average EEG response evoked by TMS. In this setting, the operator can deliver a limited number of TMS pulses (10–20) and visualize within less than a minute with unprecedented clarity the build-up, the quality, and the amplitude of TEPs. Here, the user can (*i*) ascertain whether the TEPs are characterized by features that are consistent with an EEG response to direct cortical stimulation and (*ii*) quantify the strength of the initial neuronal response. If the visualized TEPs meet the desired criteria (see the next section 2.2.1), the operator can start the measurements and data collection with a full set of trials (typically at least 100). Otherwise, she/he can further refine stimulation parameters. Below, we illustrate, using data obtained during a typical experiment, how `rt-TEP` can guide the operator in this decision process towards the maximization of the SNR of TEPs.

#### 2.2.1 Using rt-TEP during a typical experiment

We here describe a typical experiment in which `rt-TEP` is used as a guide to titrate stimulation parameters to a desired end point set by the user. While the specific equipment employed as well as the specific end point (i.e., the desired amplitude of the early TEP) may vary depending on the available set-up and the goal of the experimenter, this example is meant to illustrate the general workflow and capability of `rt-TEP`.

In this experiment, the user wants to replicate the TEPs amplitude and waveforms obtained in previous studies when stimulating the premotor cortex of healthy controls (Rosanova et al., 2009). These responses are characterized by high-amplitude (>10 μV) early (< 50□ms) components that are specific for the angle and site of stimulation. These high-amplitude TEPs are different from the responses reported by other studies in healthy subjects (see for example the comparison in Figure 1 in Belardinelli et al., 2019) and serve as a control for studies in brain-injured patients (Casarotto et al., 2016; Rosanova et al., 2018; Sarasso et al., 2021), in which similar amplitude criteria (early components >10 μV) are set to warrant high SNR for subsequent analysis. The available equipment includes neuronavigated TMS (NBS4, Nexstim Plc, Finland) and a 64-channel DC EEG amplifier (Brainamp, BrainProducts GmbH, Germany). In particular, a 62-channel EEG cap with a 10–20 montage and two EOG channels placed with a diagonal montage are connected to the amplifier. Common reference and ground electrodes are placed on the forehead, i.e., far from the coil to avoid possible injection of TMS-related artifacts into the reference. Data are recorded at a 5-kHz sampling rate with DC-to-1000-Hz hardware filtering bandwidth and with 0.5-μV amplitude resolution. Skin preparation is performed to obtain < 5kΩ impedance at all electrodes.

The operator exploits the available anatomical information to set the initial stimulation parameters; using online navigation on individual anatomical magnetic resonance images, she/he targets the premotor cortex a few centimeters left from the midline and orients the induced E-field orthogonally with respect to the underlying superior frontal gyrus. Besides anatomical information, the operator also uses physiological knowledge and initially sets stimulation intensity at 38% of the maximum stimulator output (MSO) corresponding to the resting motor threshold (rMT) previously assessed in the same subject. A masking noise is continuously played during TMS stimulation through in-ear earphones and titrated, within safety limits, in order to mask the coil click. In this case, custom software (Russo et al., 2021) generating a device-specific and subject-specific masking noise is used; at the beginning of the experiment, a few pulses are delivered while iteratively adjusting the noise parameters until the subject reports that the TMS click is not discernible.

`rt-TEP` is initialized with the IP address of the server computer (directly connected to the EEG amplifier), with the ordered set of channels streamed out by the amplifier, with the label of the TMS trigger, with the specification of the two EOG channels such that a bipolar signal is visualized, with the length of the time windows for RAW data check (i.e., 5 s) and for display of EEG responses to TMS both in single-trial and average mode (i.e., from −100 to 400 ms with respect to TMS onset time).

The operator now delivers a few test pulses in the single-trial mode to identify and control artifacts potentially affecting the early post-stimulus time interval. The pulse artifact is checked in the `rt-TEP` single-trial mode and it is masked by setting the TMS-artifact time window from −2 to +5 ms. Recharge artifact is prevented by setting the time of recharging of the stimulator’s capacitors at 900 ms after the pulse. Discharge or muscle artifacts are minimized as described in the previous section (see section 2.1).

After switching to the average mode, the operator starts a short block of 20 stimuli at a jittered inter-pulse interval of 2–2.3 s and observes, in real time, the build-up of the EEG response on the `rt-TEP` monitor. Within a minute, the operator can appreciate the resulting average TEP (**Figure 4A**) and notices that early (< 50 ms) components are absent or very small (below 2 μV) in the channels surrounding the stimulation target (black cross), indicating that the parameters based on a priori information are not effective in eliciting an immediate local response. Thus, she/he decides to increase stimulation intensity to 46%MSO (corresponding to 120% rMT) and delivers another set of 20 pulses, while keeping the same coil position and orientation. Now, the operator detects on the `rt-TEP` display two new elements (**Figure 4B**). First, early components clearly emerge immediately after the pulse at the stimulated site, though they are still small (4 μV in F1) with respect to the desired end point (10 μV). Second, she/he notices the appearance of larger (6–8 μV) negative–positive deflections between 100–200 ms. Their large amplitude with respect to the early components and their central distribution, without any visible asymmetry related to the stimulation side, suggest the presence of auditory-evoked potentials (Nikouline et al., 1999). Based on these two observations, the user decides to further increase stimulation intensity and to adjust noise masking, which was previously titrated to a lower intensity (i.e., 38% MSO).

**Figure 4.**
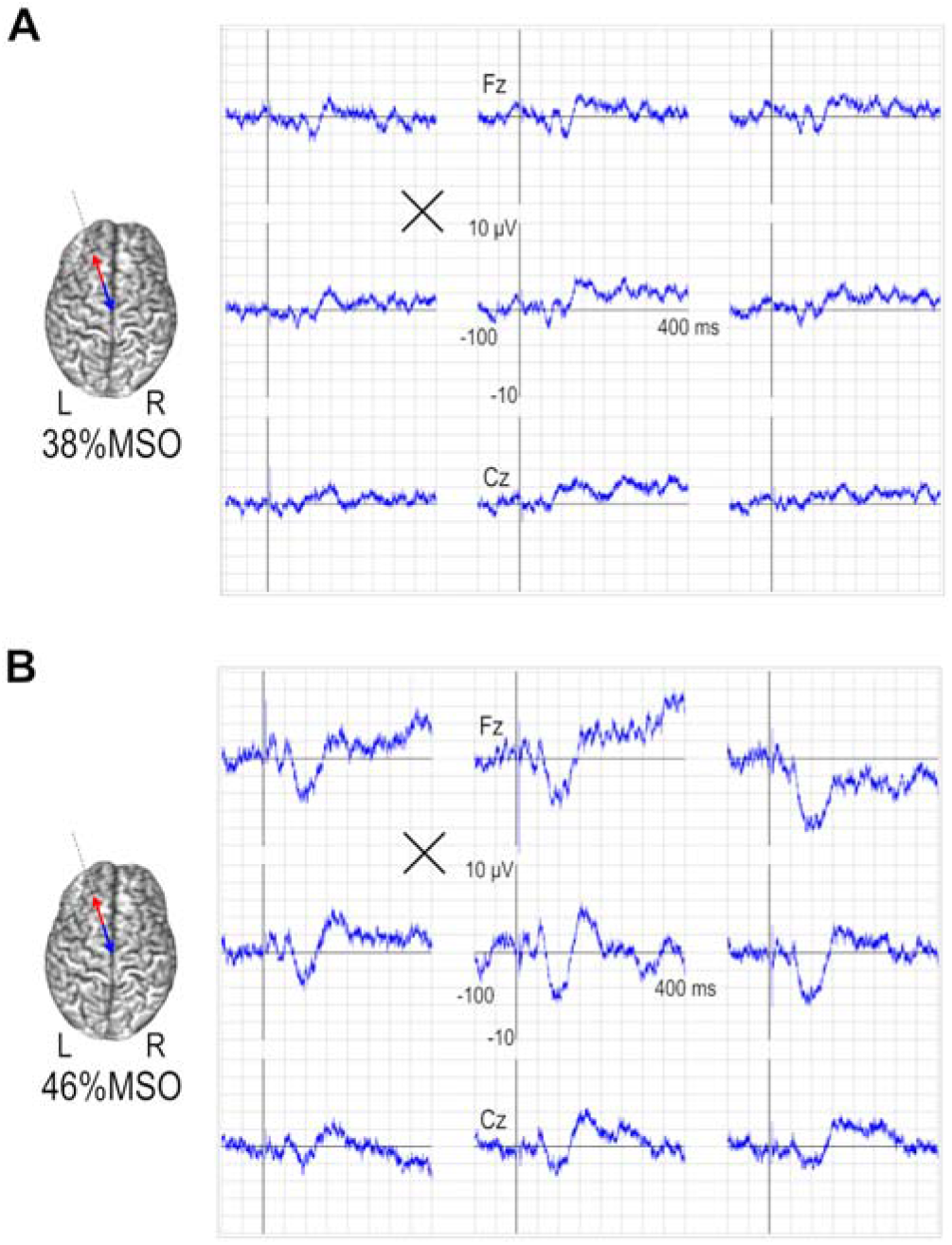
20-trial average EEG responses to TMS (zoom on frontocentral channels) when stimulating the same target on the left premotor cortex at 38% MSO (A) and 46% MSO (B). Stimulation parameters are also depicted on a rendered brain (left). EEG channels are displayed in the average reference. MSO = maximum stimulator output; L = left; R = right.

She/he thus sets the TMS intensity to 50% of the MSO and optimizes noise masking by increasing its level (always within safety limits) and/or by changing its spectral characteristics based on the subject’s report (see Russo et al., 2021).

After a new sequence of 20 pulses and another minute, the operator can appreciate the effects of the above-mentioned adjustments (**Figure 5A**); early components are larger (7 μV in F1) and are followed by the emergence of fast oscillations, whereas the auditory-like N100–P200 sequence is obliterated. Overall, the response shows an asymmetric distribution, consistent with the stimulation of a lateral target. Such real-time feedback suggests that only small adjustments are needed to attain the desired end point. To do this, the operator has different options: either further increasing TMS intensity or changing the coil angle, a modification that is known to have a significant impact on the immediate effects of TMS on the underlying circuits (Bonato et al., 2006; Casarotto et al., 2010; Tervo et al., 2021). The operator considers exploring a different angle as a first choice as this is less likely to require further adjustments of noise masking. She/he thus rotates the coil by 30° counterclockwise, while keeping the intensity constant. Delivering a new block of pulses confirms the effectiveness of this new parameter setting in achieving the desired end point (early components >10 μV) as the `rt-TEP` monitor shows a high-amplitude early response at the site of stimulation (14 μV in F1) (**Figure 5B**). The operator now sets the stimulator to deliver a sequence of 150 pulses to perform data acquisition for subsequent analysis. **Figure 5C** shows that similar results could have also been obtained, without rotating the coil, by increasing MSO by another 5%.

**Figure 5.**
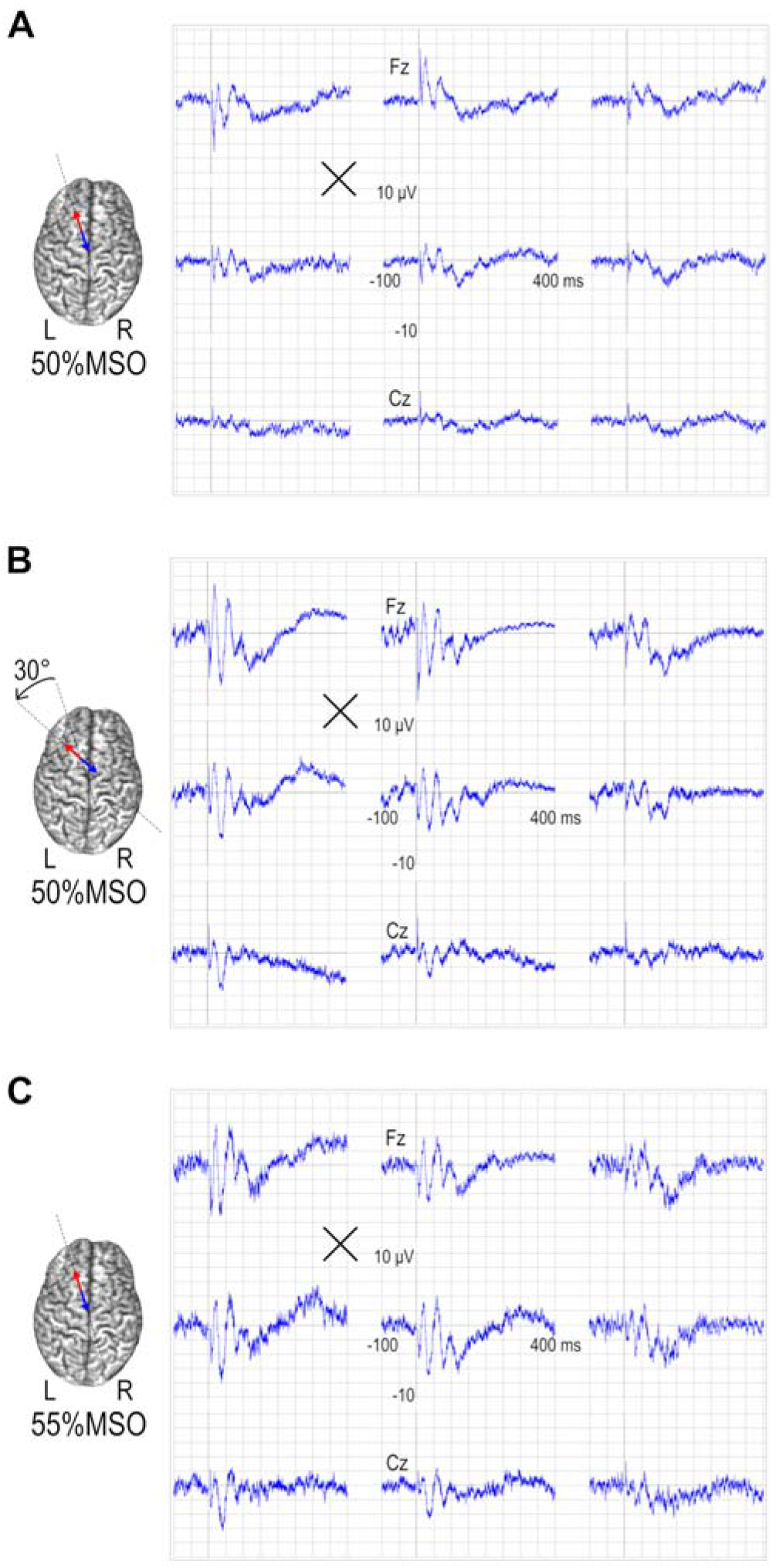
20-trial average EEG responses to TMS (zoom on frontocentral channels) when stimulating the same target on the left premotor cortex (black cross) at 50% MSO with a certain orientation (A), at 50% MSO after rotating the coil orientation by 30° counterclockwise (B) and at 55% MSO with the same coil orientation used in (A). Stimulation parameters are also depicted on a rendered brain (left). EEG channels are displayed in average reference. MSO = maximum stimulator output; L = left; R = right.

**Figure 6** shows the TEPs obtained after averaging 20 trials and at the end of the session after averaging 150 trials (same parameters as in **Figure 5B**). Here, one can note two important aspects. First, the amplitudes of the early components observed during the parameter search (average of 20 trials – **Figure 6A**) provide a good estimation of their final value (average of 150 trials – **Figure 6B**). Second, the quality of the acquired TEPs can be readily appreciated by the user at the end of the experiment without the need for any pre-processing (e.g., detrending, independent component analysis, …).

**Figure 6.**
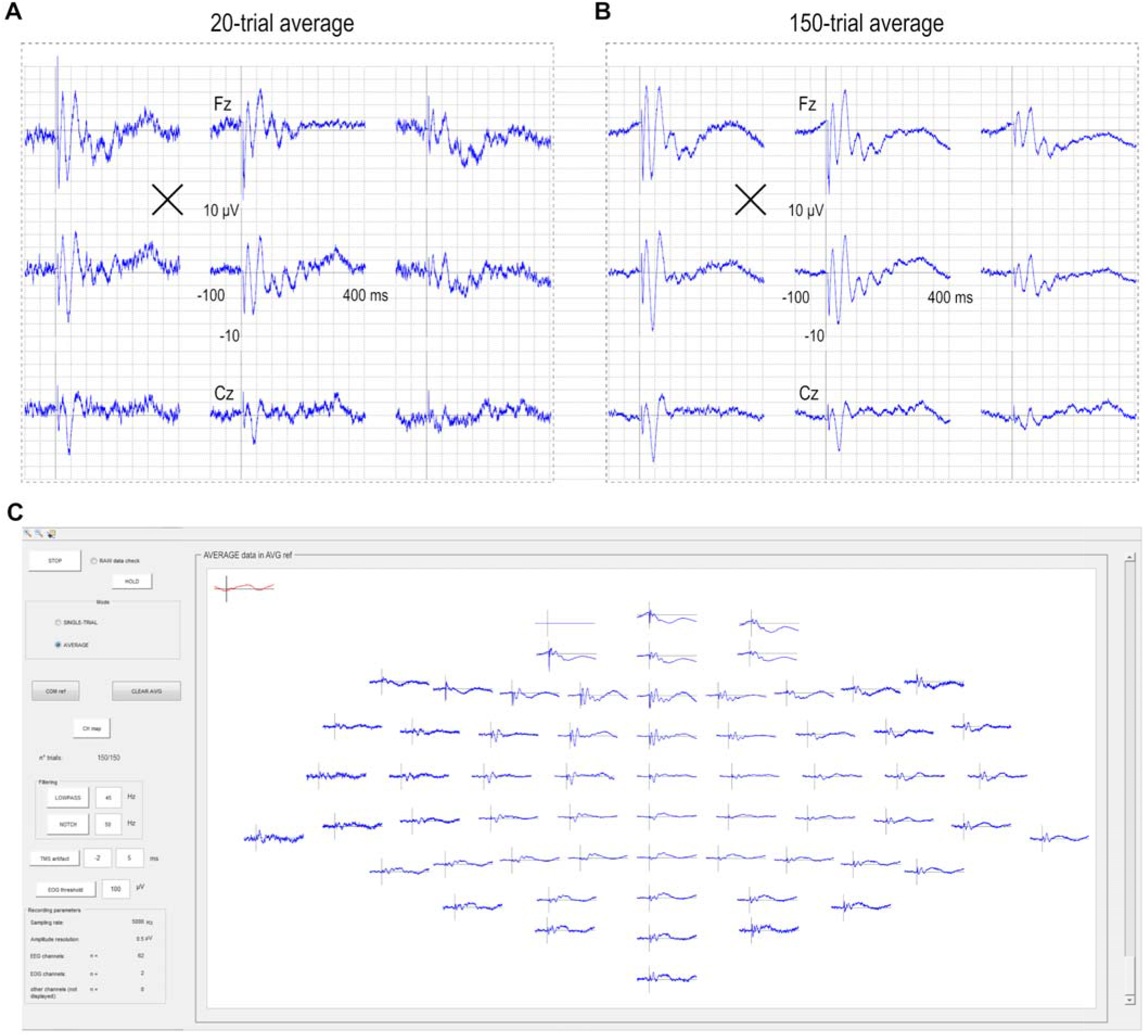
EEG responses to TMS when stimulating the left premotor cortex (black cross) at 50% MSO with a certain orientation (as in Figure 5B) after averaging 20 trials (A for a zoom on frontocentral channels) and 150 trials (B for a zoom on frontocentral channels, C for the all-channels display). EEG channels (blue trace) are displayed in average reference whereas EOG (red trace) is displayed in bipolar montage. MSO = maximum stimulator output.

The importance of setting stimulation parameters before starting the actual measurement, in order to optimize the initial cortical activation and to minimize obvious artifacts, is further illustrated in **Figure 7**. This figure compares directly the final average TEPs (150 trials) collected during three sessions (corresponding, from top to bottom to the stimulation parameters set in **Figure 4A**, **Figure 4B** and **Figure 5B**). Although all these responses have been obtained by setting stimulation parameters based on reasonable a priori anatomical (position and orientation on the cortical gyrus) and physiological (%MSO at or above rMT) assumptions, they differ in fundamental ways. The responses in **Figures 7A** and **7B** show absent/small early activations and are characterized by larger, late symmetric components which are maximal over midline channels. These waveforms are hardly consistent with the effects of direct cortical stimulation, which is expected to trigger responses that are large immediately after the pulse and specific for the stimulation site (Keller et al., 2014; Kundu et al., 2020). Conversely, the TEP reported in **Figure 7C** is similar to those reported in previous studies (Rosanova et al., 2009; Casarotto et al., 2016; Sinitsyn et al., 2020) and fulfils these basic criteria. In this case, a strong initial activation is followed by an overall asymmetric wave shape with high SNR. As described above, reproducing this kind of responses only required maximising the immediate impact of TMS on early (< 50 ms) components through slight adjustments of the intensity (5-10% MSO) and/or the angle of stimulation, while at the same time optimizing noise masking. Making such adjustments is relatively straightforward but would be impossible based on a priori information alone and can only be done if the operator is guided in real-time by an informative feedback about the immediate effects of TMS, such as the one provided by `rt-TEP`.

**Figure 7.**
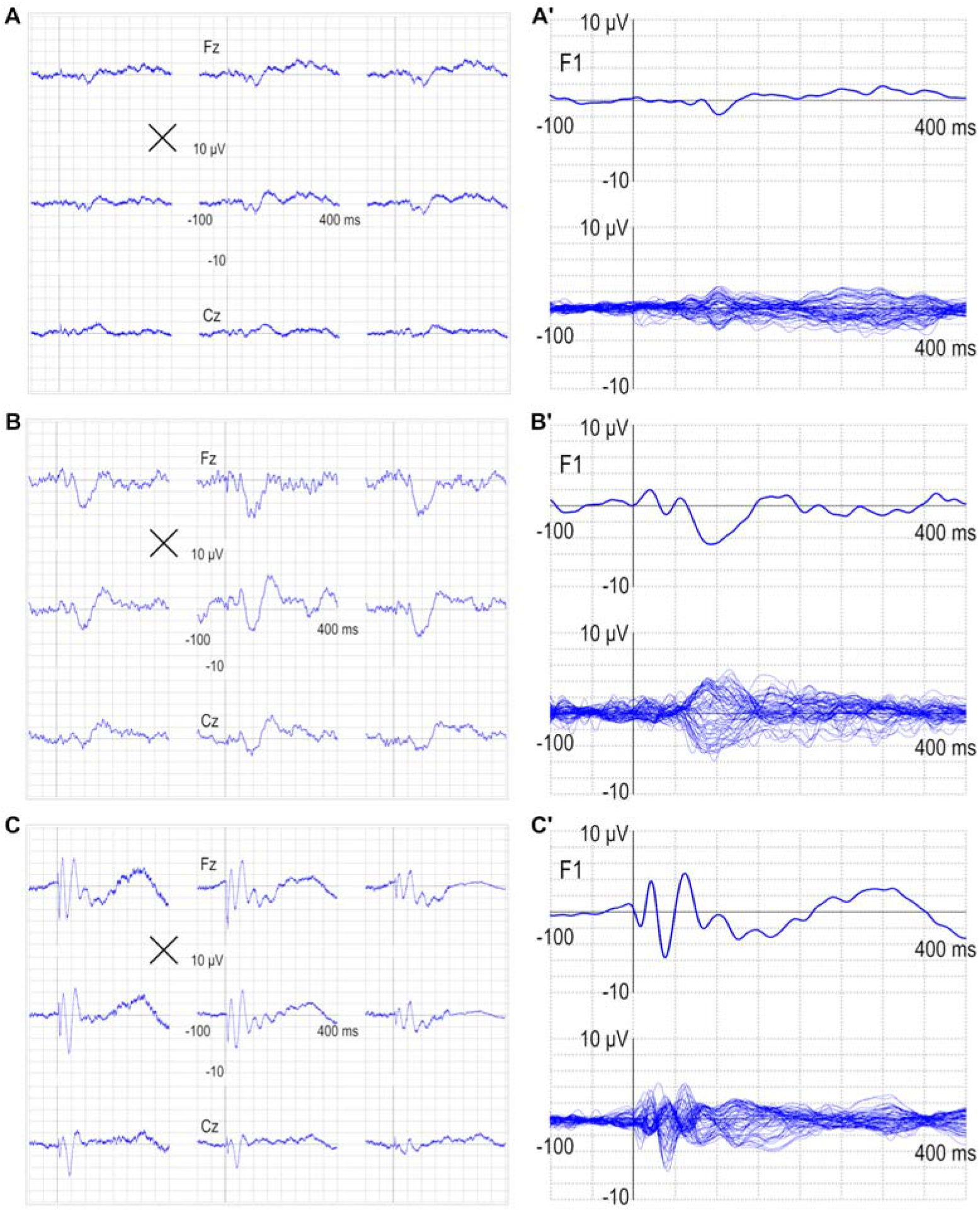
150-trial average EEG responses to TMS (zoom on frontocentral channels) when stimulating the same target on the left premotor cortex (black cross) at 38% MSO (A), at 46% MSO (B) and at 50% MSO with adjusted noise masking after rotating the coil orientation by 30° counterclockwise (C). See **Figures 4A, 4B** and **5B** for the corresponding 20-trial averages. EEG channels are displayed in the average reference. The corresponding F1 channel and butterfly plot (panels A’, B’ and C’) have been low-pass filtered at 45 Hz. MSO = maximum stimulator output.

## 3. Discussion

With this paper, we introduce and release a novel tool to facilitate the acquisition of TEPs based on a real-time readout of the immediate impact of TMS on the underlying neuronal circuits. In short, `rt-TEP` guides the user through two sequential steps. In the first step (i.e., single-trial mode), artifacts potentially contaminating the early post-stimulus period are abolished or minimized. In the second step (average mode), stimulation parameters are adjusted until reaching the desired level of cortical activation, as judged by the amplitude of early (< 50 ms) components located under the coil. In the experiment described in the present paper, we aimed at a peak-to-peak amplitude larger than 10 μV, an endpoint that has been used in previous studies to warrant high SNR for subsequent TEP analysis in healthy and brain-injured individuals (Casarotto et al., 2016; Rosanova et al., 2018; Sinitsyn et al., 2020). While the specific endpoint can be set based on more sophisticated criteria (see **Supplementary results**) and may vary upon the experimenter’s needs and purposes, we here discuss the merit of the general approach.

### 3.1 rt-TEP: rationale and applications

As we have recalled in the Introduction, the actual impact of the TMS-induced E-field on cortical neurons is very hard to predict based on a priori information. This problem is particularly relevant when targeting cortical sites outside of the primary motor area for which the immediate readout represented by the motor evoked potential (MEP) is not available. This lack of control on stimulation effectiveness may explain a significant portion of the large variability of TEP waveforms reported in the current literature. For example, while in many TMS–EEG studies TEPs are small in amplitude, symmetric and not specific for the cortical target, in others the stimulation of the same areas gives rise to very different responses that are much larger (up to one order of magnitude), asymmetric and specific for the stimulated cortical site. As exemplified in **Figure 7**, profoundly different responses can all be obtained within the envelope of reasonable parameter settings based on a priori information, such as individual anatomy and individual rMT. In fact, this variability (including instances in which no responses to TMS are detectable) likely reflects key factors that remain unaccounted for, including microscale axon orientation, cytoarchitectonic and local input–output properties of neurons.

The idea behind `rt-TEP` is that, while these factors remain hard to predict, it is possible to bypass them by controlling and standardizing stimulation parameters based on a post-hoc readout, i.e., the amplitude of early EEG components. In this perspective, `rt-TEP` extends to any cortical area the same logic that normally applies to the primary motor cortex, which involves an adjustment of stimulation intensity to elicit MEPs with a desired amplitude. There are, however, specific challenges characterizing this EEG-based approach as compared to the classic MEP-based approach. First, early TMS-evoked EEG responses are harder to visualize than peripheral TMS-evoked EMG responses, because they can be contaminated by various types of artifacts. Third, the actual readout is represented by a spatiotemporal distribution of average potentials rather than by single-pulse EMG wave, which require a dedicated visualization mode (i.e., average reference, topographic arrangement) to appreciate the spatial properties and the amplitude of the brain response. The key function of `rt-TEP` is to assist the operator in overcoming these problems in order to visualize and control the initial impact of TMS. Once the initial cortical input is effective and standardized (the independent variable), the experimenter can measure and analyse the evolution of the ensuing brain response in time and space (the dependent variable).

For example, it is possible to appreciate the specificity of TMS-evoked potentials across different stimulation sites (Casarotto et al., 2010), to analyze the frequency content of the overall EEG response to the initial perturbation (i.e., the natural frequency; Rosanova et al., 2009), and how it may be altered in pathological conditions (Ferrarelli et al., 2008, 2012; Canali et al., 2015). High signal-to-noise TEPs generated by effective perturbations can also be analyzed to derive brain complexity measures that are clinically useful to stratify patients with disorders of consciousness (Casali et al., 2013; Casarotto et al., 2016; Bodart et al., 2017; Sinitsyn et al., 2020). Likewise, ensuring effective stimulation is a key prerequisite to compare the electrophysiological properties of the stroke perilesional area to the ones of the contralateral site (Sarasso et al., 2020). Finally, once clearly visible, early components can be considered themselves as a dependent variable when assessing changes in cortical excitability in repeated measurements within individuals. Indeed, once the selected stimulation parameters (site, intensity and angle) are kept fixed with the aid of a reliable TMS-navigation system, the shape and amplitude of these components are highly reproducible across repeated studies (Casarotto et al., 2010) and are sensitive to manipulations such as electroconvulsive therapy (Casarotto et al., 2013), sleep deprivation (Chellappa et al., 2016; Gaggioni et al., 2019, 2021; Huber et al., 2013; Ly et al., 2016) and transcranial direct current stimulation (Romero Lauro et al., 2014). More generally, making sure that the initial effects of TMS are present and standardized within a given range is key to fostering the reproducibility and exchange of TMS–EEG data across laboratories.

### 3.2 rt-TEP: caveats and perspectives

A major limitation of TMS-EEG-based approaches in general is represented by unavoidable artifacts that can be present when stimulating cortical areas located directly below scalp muscle insertions (Mutanen et al., 2013). The parameter search in the single-trial mode is reliably successful in avoiding such artifacts when placing the coil over a large portion of the head; in our experience, a clean shot on early components can be easily obtained in correspondence of an area of the cortex spanning from Brodmann’s area (BA) 18/19 to BA 6/9 and extending a few centimeters around the midline, including the hand motor areas (Rosanova et al., 2009; Casali et al., 2010; Ferrarelli et al., 2012; Fecchio et al., 2017). The likelihood of success decreases markedly as the coil is moved anteriorly to dorsolateral prefrontal cortex (DLPFC) or posteriorly to BA 17 and is virtually null on language areas.

Finally, the ultimate success of the overall `rt-TEP` procedure depends on the available hardware. For example, active amplifiers tend to induce early discharge artifacts that are more prominent and difficult to eliminate, often masking early components. Also, the accuracy of the TMS-navigation unit at hand is a key factor; indeed, the settings (coil position coordinates and rotation) identified during the parameter search in `rt-TEP` must be precisely retrieved and held steady throughout the actual measurement. Finally, TMS hardware, coils and pulse waveshapes can differ largely in their focality, efficacy on cortical circuits and collateral effects (magnetic artifacts, scalp and auditory stimulation) (Koponen et al., 2020; Van Doren et al., 2015). As major theoretical and technical efforts are currently ongoing to optimize these factors, `rt-TEP` may represent a useful tool to empirically explore and compare the effectiveness of the different solutions.

As described in this paper, in order to achieve effective stimulation, the adjustment of TMS parameters may involve, in addition to intensity changes, small coil rotations. Although a few manual rotations are generally effective in increasing early TEPs, a systematic search of the optimal E-field orientation is practically unfeasible. Such fine tuning requires more sophisticated strategies and hardware, such as an EEG-based adaptive search algorithm coupled with an electronically-controlled two-coil transducer (Souza et al., 2021; Tervo et al., 2020, 2021). Combining `rt-TEP` with advanced closed-loop systems represents a promising strategy whereby fundamental stimulation parameters are first set by the operator based on visual feed-back and then automatically optimized in a closed-loop fashion.

### 3.3 rt-TEP: why use it?

The tool presented in this paper offers an informative sequence of simple visualization modes to facilitate the successful acquisition of TEPs that is not currently implemented in any commercial software. Below, we would like to highlight key reasons that should motivate the incorporation of `rt-TEP` in future experimental designs.

First and foremost, researchers and clinicians alike would like to avoid situations such as that illustrated in **Figure 7A** where, in spite of reasonable a priori assumptions, TMS has no or little impact on the underlying cortex. Such occurrences, which represent a clear drawback not only for TMS–EEG studies but also for interventional protocols (plasticity), can be readily controlled for and prevented with `rt-TEP`.

Second, as already mentioned, standardizing the initial effects of TMS within a given range may mitigate the problem of reproducibility currently affecting the TMS–EEG literature (Belardinelli et al., 2019).

Third, `rt-TEP` allows increasing the cortical impact of TMS in ways (small coil translation and rotations) that do not necessarily involve, or minimize, the need for large increases of stimulation intensity (**Figure 5B**). Such optimization of the effects of genuine cortical activation over the collateral sensory effects of the coil’s discharge is key to improve the quality and informativeness of TEPs.

Fourth, through a series of visualization steps, `rt-TEP` guides the operator in eliminating/minimizing obvious artifacts and confounding factors during data collection, including muscle twitches and auditory evoked potentials, which must be otherwise eliminated during post processing. This is an important advantage, in view of the strong sensory input associated with muscle twitches and considering the limitations inherent to many off-line artifact rejection algorithms (such as principal component analysis - PCA and independent component analysis - ICA) (Biabani et al., 2019; Belardinelli et al., 2019; Bertazzoli et al., 2021).

Finally, `rt-TEP` provides the experimenter with clear, real-time feedback about data quality already during the experiment (**Figure 6**). This prevents discovering poor TEP quality only at a later stage, which is particularly problematic when a second measurement is not an option, as it is often the case in patients.

More generally, real-time feedback about the effect of TMS on the underlying tissue may render TMS–EEG more reliable and akin to other measuring tools that have proven to be extremely powerful in medicine. As exemplified in this paper, the TMS probe, just like an ultrasound probe, requires informed handling in order to recover a strong signal of interest amidst layers of noise. In daily practice, echography operators are involved in a similar task; they orient the probe until they recover on their monitors a robust echo from the target structure; when basic SNR criteria are met, only then can the actual measurement start. Thanks to an effective real-time readout and with some practice, these operator-dependent procedures become second nature and standardized to the point of supporting important medical decisions. By comparison, TMS–EEG is still in its infancy, but we hope that the release of `rt-TEP` software may represent a step further in this direction.

## Notes

### Competing Interest Statement

The authors have declared no competing interest.

### Summary of Updates

Title has been slightly modified. A small but important correction has been made to paragraph "2.1.4 Discharge artifacts": the artifact produced by discharge of the skin-related capacitors is actually more complex than and exponential decay (Freche et al., 2018). In the original sentence "non-" was missing by mistake. Thus, the revised sentence now read as follows: "Immediately after being charged by the pulse, these capacitances discharge and produce a non-exponentially decaying artifact on the recorded signal, which can last several tens of ms."

