## Supplementary material for "The rt-TEP tool: real-time visualization of TMS-Evoked Potential to maximize cortical activation and minimize artifacts" (from another version of this preprint: version 2, 2021-09-21)

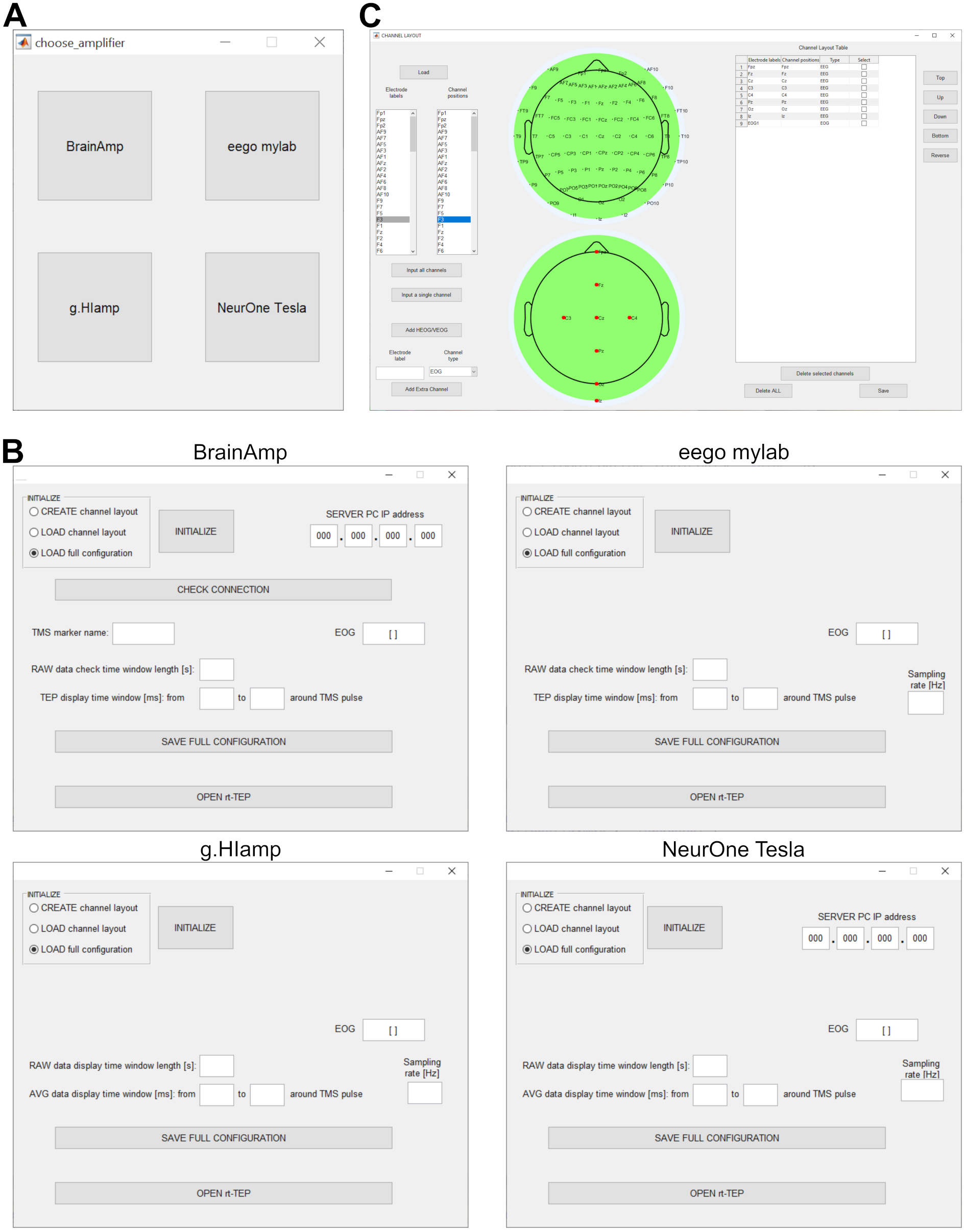

**Supplementary Figure 1.** The opening window of rt-TEP (A) allows selecting the EEG amplifier in use. Then, configuration parameters must be indicated within an amplifier-specific initialization window (B) before starting data display in real-time. When selecting the *CREATE channel layout* radio button and pressing INITIALIZE, a dedicated interface appears (C) to create the channel layout currently in use by picking and ordering EEG channels from a comprehensive list provided by the software.

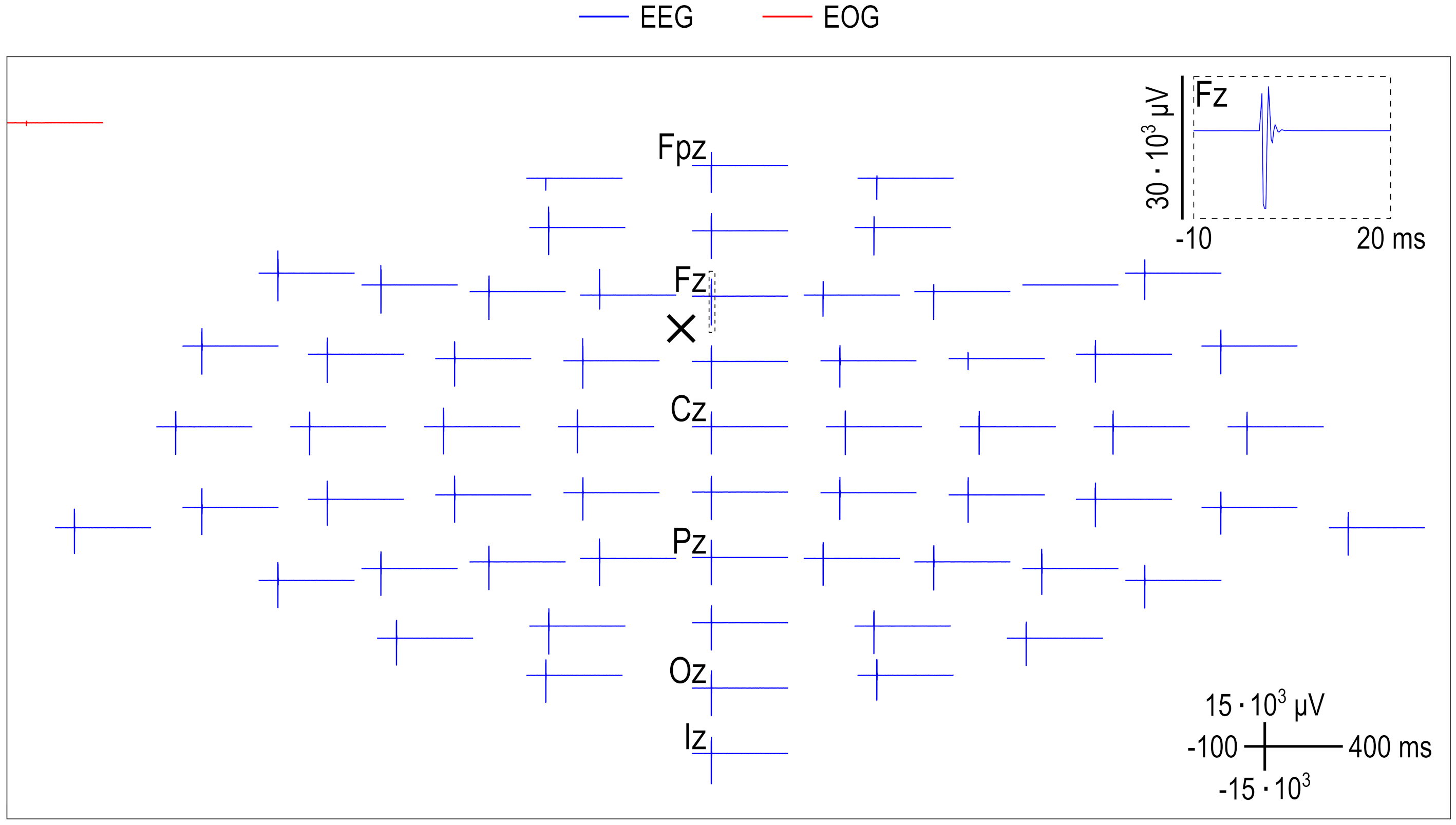

**Supplementary Figure 2.** Example of a pulse artifact on single-trial data: EEG channels (blue traces) are displayed in common physical reference, whereas EOG (red trace) is displayed in bipolar montage. A zoom-in of the measured signal in channel Fz around the TMS pulse is shown in the top right inset. The EEG was recorded with a BrainAmp DC amplifier and TMS was delivered with an NBS4 Nexstim system at 50% of its maximum stimulator output with the figure-of-eight coil center positioned at the location corresponding to the black cross.

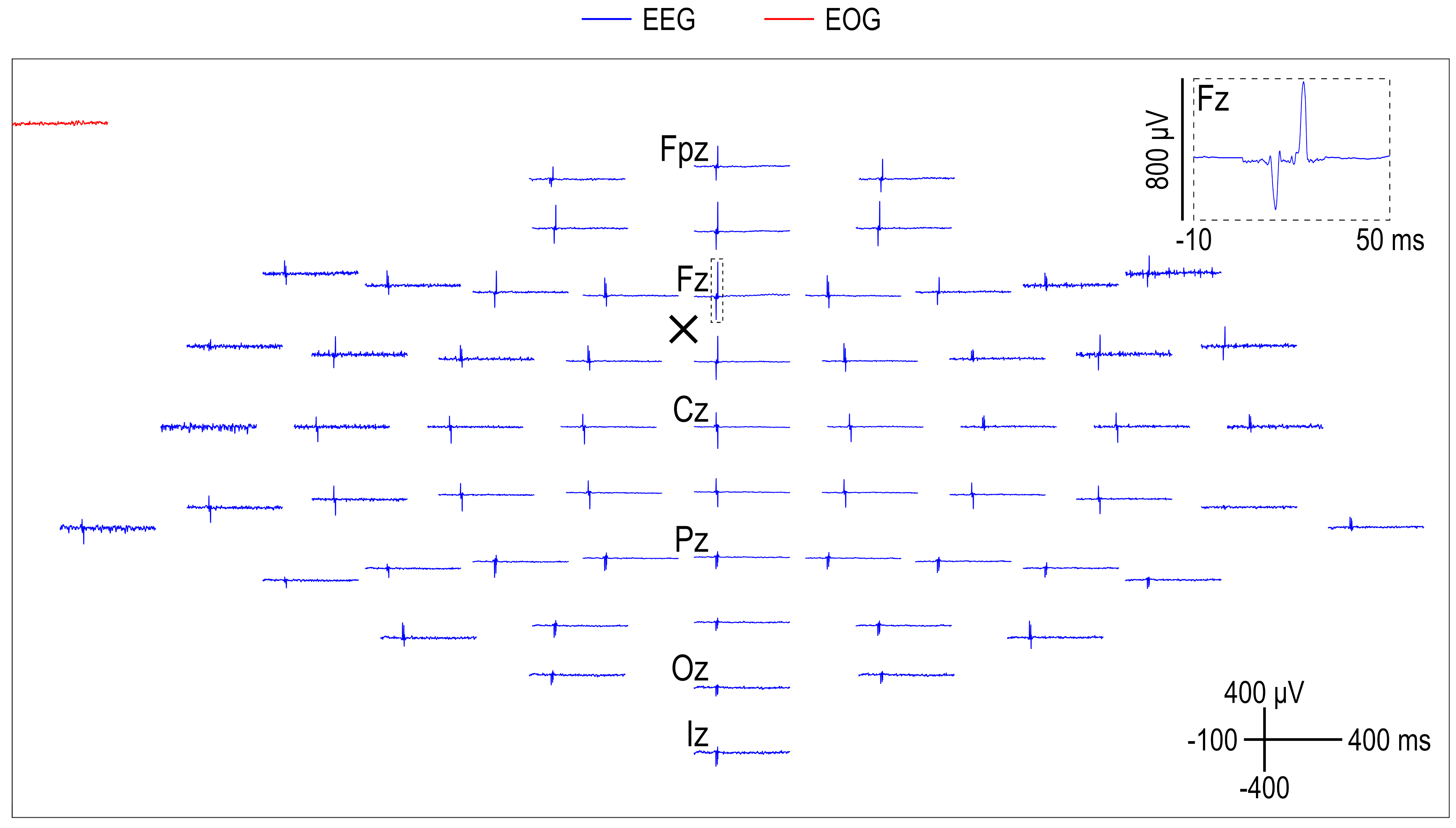

**Supplementary Figure 3.** Example of a recharge artifact on single-trial data: EEG channels (blue traces) are displayed in average reference, whereas EOG (red trace) is displayed in bipolar montage. A zoom-in of channel Fz around the TMS pulse is shown in the top right inset. Data were recorded with a BrainAmp DC amplifier and TMS was delivered with a Rapid^2^ stimulator (Magstim Ltd, UK) equipped with a D70 Remote Coil at 50% of its maximum stimulator output with the figure-of-eight coil center positioned at the location corresponding to the black cross.

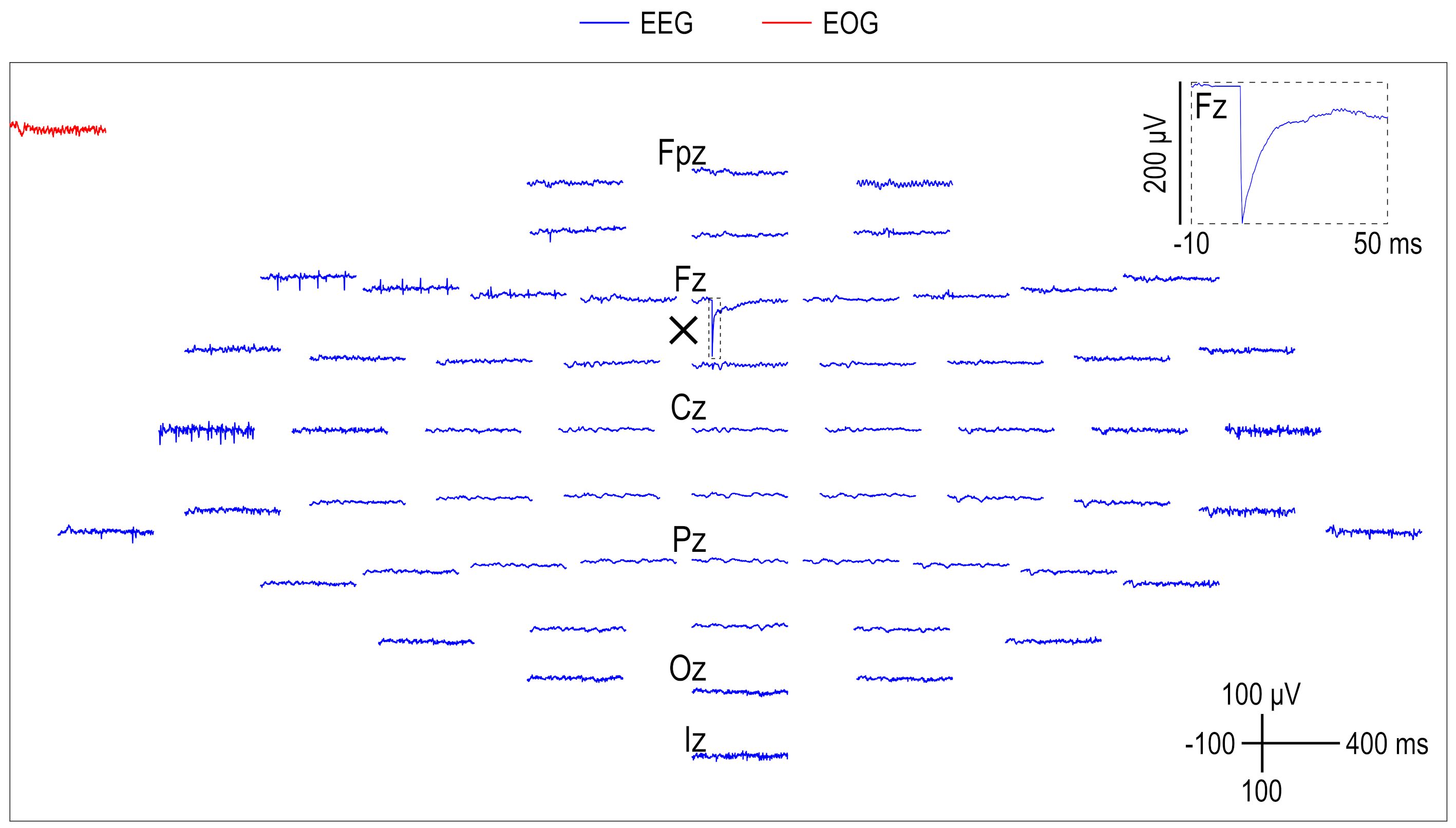

**Supplementary Figure 4.** Example of a discharge artifact on single-trial data: EEG channels (blue traces) are displayed in average reference, whereas EOG (red trace) is displayed in bipolar montage. A zoom-in of channel Fz around the TMS pulse is shown in the top right inset. Data were recorded with a BrainAmp DC amplifier and TMS was delivered with an NBS4 Nexstim system at 50% of its maximum stimulator output with the figure-of-eight coil center positioned at the location corresponding to the black cross.

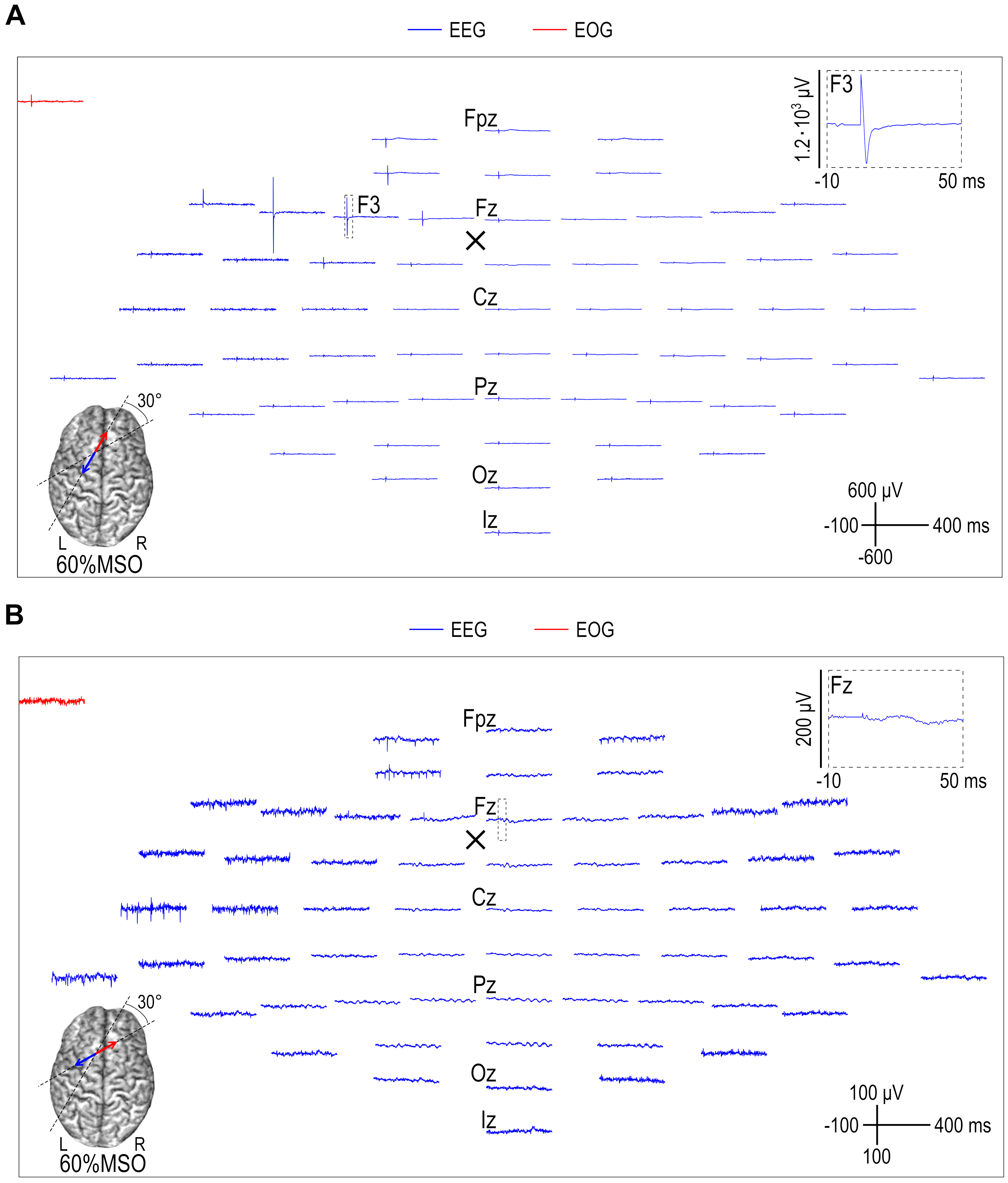

**Supplementary Figure 5.** Example of muscle artifact on single-trial data. TMS was applied on the left frontal cortex (black cross) at 60% of MSO: the orientation of the induced E-field differs by 30° between panels A and B, as depicted on a rendered brain. The pulse artifact was visually masked by flattening the signal between –2 and +5 ms. EEG channels (blue traces) are displayed in the average reference, whereas EOG (red trace) is displayed in bipolar reference. EEG data were collected with a BrainAmp DC amplifier and TMS was delivered with an NBS4 Nexstim system. MSO = maximum stimulator output; L = left; R = right.

### **Supplementary methods**

Data reported in this paper were collected with a 64-channel DC EEG amplifier (Brainamp, BrainProducts GmbH, Germany), including 62 EEG channels (10–20 montage) and two EOG channels (diagonal montage). Two additional electrodes were placed on the forehead and used as common physical reference and ground. Data were recorded at 5-kHz sampling rate with DC-to-1000-Hz hardware filtering bandwidth and with 0.5-µV amplitude resolution. Unless otherwise specified (see **Supplementary Figure 2**), TMS has been delivered with an NBS4 system (Nexstim Plc, Finland), which includes a neuronavigation software and is equipped with a figure-of-eight coil (mean/outer winding diameter 50/70 mm, biphasic pulse of length 230 µs, focal area of the stimulation 0.68 cm^2^). The induced E-field depicted in **Figures 3–5** and **Supplementary Figure 5** on a rendered brain is represented by two opposite arrows: the intersection between the two arrows indicates the TMS target, while the direction of the induced E-field on the cortex first corresponds to the red arrow and then to the blue arrow after reversing the current flow during biphasic pulse delivery.

### **Supplementary results**

From a *quantitative perspective*, average EEG potentials evoked in the first 50 ms in the channel closest to the stimulation site should be large enough to emerge from spontaneous EEG activity. Using simulated data, it is possible to estimate what is the minimum amplitude of the single-trial evoked response that can be almost perfectly recovered from baseline EEG after averaging a certain number of trials: then, the corresponding peak-to-peak amplitude that should be measured after averaging a limited number of trials can be inferred in order to predict in advance whether TEPs have a high signal-to-noise ratio.

We considered a database of TMS–EEG recordings in 30 healthy awake subjects: in particular, we selected the channels {AF3, AFz, AF4, F1, Fz, F2, FC1, FCz, FC2, C1, Cz, C2} that are usually close to the stimulation site when TMS is targeted over the premotor cortex, either on the left or right hemisphere. From each trial of the database in each channel, we extracted a 150-ms-long segment of spontaneous EEG from the baseline period (**Supplementary Figure 6A**). Then, we considered the EEG responses to TMS of the left premotor cortex freely available from (Belardinelli et al., 2019): in particular, we used the first 50 ms of the TMS-evoked potential measured in channel Fz (in average reference and normalized to 1 µV peak-to-peak amplitude) as a template, preceded by 100 ms of flat baseline (**Supplementary Figure 6B**). We summed the template multiplied by a specific weight (β) to the spontaneous EEG segments (simulated trials; **Supplementary Figure 6C**): β was iteratively estimated until the statistical fit (R^2^) between the weighted template and the average of *n*=80 simulated trials was > 0.95. Finally, the minimum desirable amplitude of TMS-evoked potentials was computed as the peak-to-peak amplitude (V_pp_) of a 20-trial average between the known latencies of the positive and negative peaks of the template. To limit the effects of possible outlier trials, for each channel of the database, we computed 1000 surrogate averages by randomly selecting 20 trials from the first 80 trials of the recording and then we reported the mean measurement across surrogates. **Supplementary Figure 6D** shows the percentage distribution of V_pp_ values estimated from 360 statistical fits (30 subjects × 12 channels): the minimum amplitude was larger than 4.3 µV in 90% of the cases, with a median value of 7.3 µV (5.3–10.9 µV interquartile range). The signal-to-noise ratio (computed as the ratio between RMS[0…50 ms] and RMS[–100…0 ms]; RMS = root-mean-square level) of the 80-trial average was larger than 2.9 in 90% of the cases (median value 4.8) (REF Parks). Finally, the minimum desirable amplitude of TMS-evoked potentials after 20 trials (V_pp_) was significantly correlated (*p* < 0.0001) with the RMS[–100…0 ms] (**Supplementary Figure 6E**).

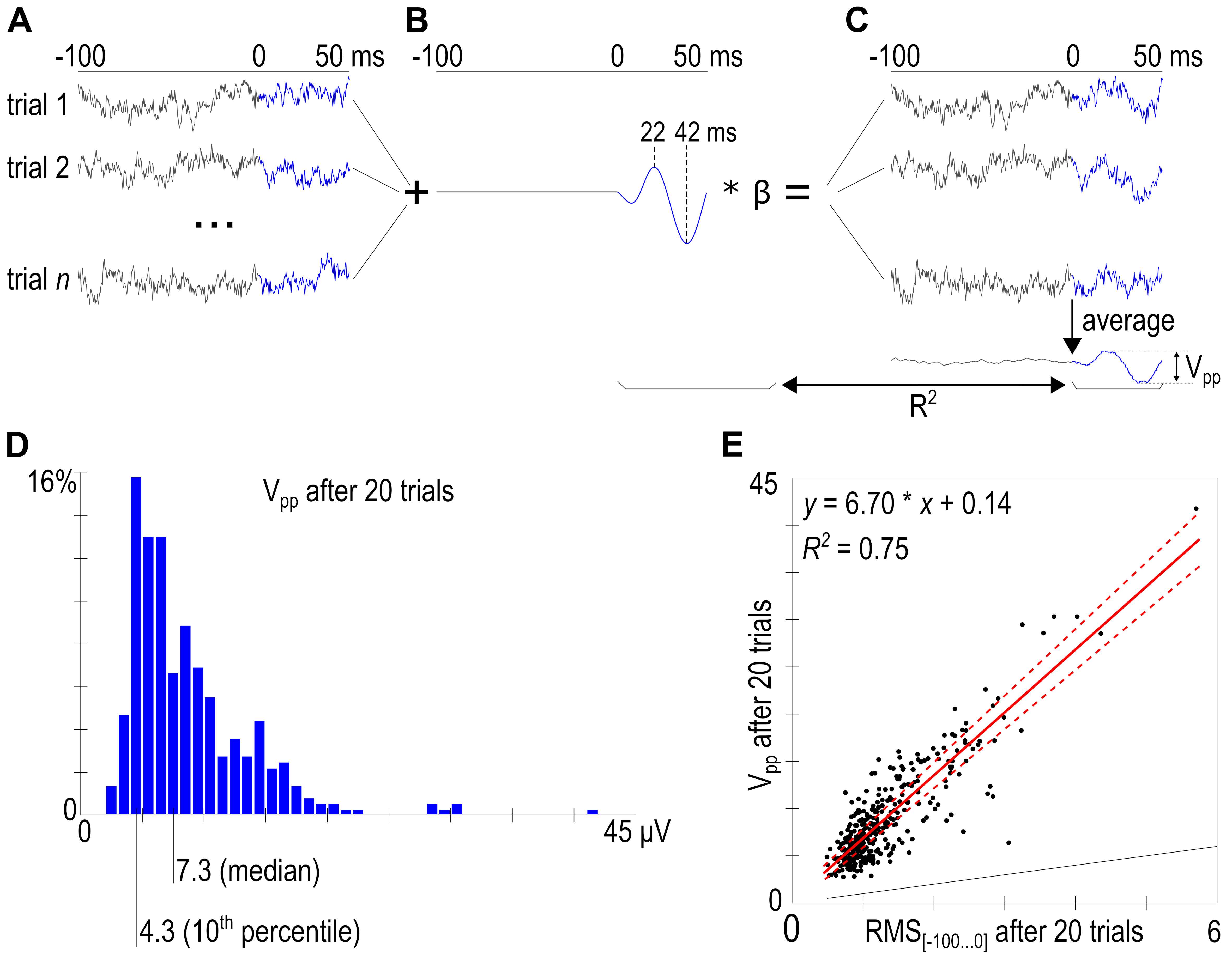

**Supplementary Figure 6.** Outline of the mathematical procedure to estimate what is the minimum amplitude of the single-trial evoked response that can be almost perfectly recovered from baseline EEG after averaging a certain number of trials. Starting from a database of TMS–EEG recordings in 30 healthy awake subjects, for each trial (out of *n*) in each channel (among {AF3, AFz, AF4, F1, Fz, F2, FC1, FCz, FC2, C1, Cz, C2}), 150-ms-long segments of spontaneous EEG (A) are summed to a TEP template (B) (taken from Belardinelli et al., 2019) multiplied by a specific weight (β), thus obtaining a surrogate signal (C). β is iteratively estimated until the statistical fit (R^2^) between the weighted template and the average of *n*=80 simulated trials is > 0.95. The minimum desirable amplitude of TMS-evoked potentials is computed as the peak-to-peak amplitude (V_pp_) of a 20-trial average between the known latencies of the positive and negative peaks of the template. The percentage distribution of V_pp_ values estimated from 360 statistical fits (30 subjects × 12 channels) is reported in panel D. V_pp_ is significantly correlated (*p* < 0.0001) with the RMS[–100…0 ms], as shown in panel E.
